# Knockdown of the TRPM4 channel alters cardiac electrophysiology and hemodynamics in a sex- and age-dependent manner in mice

**DOI:** 10.1101/2022.10.26.513825

**Authors:** Prakash Arullampalam, Maria C. Essers, Jean-Sébastien Rougier, Hugues Abriel

## Abstract

**Background:** TRPM4 is a calcium-activated, voltage-modulated, non-selective ion channel widely expressed in various types of cells and tissues. TRPM4 regulates the influx of sodium ions, thus playing a role in regulating the membrane potential. In the heart, TRPM4 is expressed in both cardiomyocytes and cells of the conductive pathways. Clinical studies have linked *TRPM4* mutations to several cardiac disorders. While data from experimental studies have demonstrated TRPM4’s functional significance in cardiac physiology, its exact roles in the heart remain unclear.

**Aim:** To investigate the role of TRPM4 in cardiac physiology in a newly generated knockdown *Trpm4* mouse model.

**Methods and results:** Male and female *Trpm4* knockdown (*Trpm4* ^-/-^) and wild-type mice 5- to 12-weeks-old (young) or 24-week-old or more (adult) were characterized using a multimodal approach, encompassing surface electrocardiograms (ECG), echocardiography recordings, pseudo and intracardiac ECGs, western blots, and mRNA quantifications. The assessment of cardiac electrophysiology by surface ECGs revealed no significant differences between wild type and *Trpm4* ^-/-^ 5- to 12-weeks-old mice of either sex. Above 24 weeks of age, adult male *Trpm4* ^-/-^ mice showed significantly reduced heart rate and increased heart rate variability. Echocardiography reveals that only adult male *Trpm4* ^-/-^ mice exhibited slight left ventricular hypertrophic alterations with an alteration of the mitral valve pressure half time, the mitral valve E/A ratio, the isovolumetric relaxation time, and the mitral valve deceleration. In addition, an assessment of the right ventricular systolic function by scanning the pulmonary valve highlighted an alteration in pulmonary valve peak velocity and pressure in male *Trpm4* ^-/-^ adult mice. Finally, intracardiac ECG recordings showed that the application of 5 µM NBA triggered a third-degree atrioventricular block on 40% of wild-type hearts only.

**Conclusions:** These results confirm the important role of TRPM4 in the proper structure and electrical function of the heart. It also reveals significant differences between male and female animals that have never been reported before. In addition, the investigation of the effects of NBA on heart function highlights the role of TRPM4 in atrioventricular conduction and provides the first evidence showing the efficacy of this compound on native cardiac tissues.

## INTRODUCTION

Transient receptor potential melastatin-related 4 (TRPM4) is a nonselective, cation-permeable channel modulated by transmembrane voltage (Ehara et al. 1988, Launay et al. 2002, Nilius et al. 2003). It is encoded by the mouse *Trpm4* and human *TRPM4* genes, found on chromosome 7 in mice and on chromosome 19 in humans, respectively (Guinamard et al. 2010, Launay, Fleig, Perraud, Scharenberg, Penner and Kinet 2002, Nilius et al. 2004). TRPM4 channels are widely expressed in various cells and organs, including pancreatic β-islet cells, neurons, immune cells, smooth muscle cells in blood vessels, prostate, bladder, and heart (Demion et al. 2007, Kruse et al. 2009, Partridge and Swandulla 1988, Siemen 1993).

Its conductance was first described in 1981 by Colquhoun *et al*. in rat cardiomyocytes (Colquhoun et al. 1981, Nilius and Flockerzi 2014). TRPM4 is permeable for monovalent cations Na^+^ > K^+^ >> Cs^+^ > Li^+^ and it is activated by intracellular Ca^2+^ (Launay, Fleig, Perraud, Scharenberg, Penner and Kinet 2002, Nilius, Prenen, Droogmans, Voets, Vennekens, Freichel, Wissenbach and Flockerzi 2003). Allowing the influx of sodium ions, TRPM4 activation depolarizes the cell membrane (Nilius and Flockerzi 2014). The activation of TRPM4 is modulated by PKC phosphorylation, calmodulin, and PIP_2_ (Chu and Stefani 1991, Nilius et al. 2005). Intracellular nucleotides such as ATP, ADP, AMP, and the nonhydrolyzable ATP analog adenylyl-imidodiphosphate (AMP-PNP) have been shown to inhibit the activity of the TRPM4 channel (Nilius, Prenen, Voets and Droogmans 2004).

TRPM4 dysfunction has been linked to several cardiac conduction disorders. In 2009, a genetic study linked a human *TRPM4* mutation with cardiac bundle branch block (Kruse, Schulze-Bahr, Corfield, Beckmann, Stallmeyer, Kurtbay, Ohmert, Brink and Pongs 2009). Since then, several clinical and experimental studies have linked *TRPM4* mutations with other conduction disorders such as Brugada syndrome, atrioventricular block, and right bundle branch block (Liu et al. 2013, Liu et al. 2010, Stallmeyer et al. 2012). Interestingly, both gain- and loss-of-function mutations have been suggested to lead to similar cardiac dysfunctions (Abriel et al. 2012, Demion et al. 2014). In experimental mouse studies, the cardiac action potentials from *Trpm4* ^-/-^ atrial cardiomyocytes were 20% shorter than their wild-type (WT) counterparts and suggest that TRPM4 regulates the sinus rhythm in the sinoatrial node by influencing the diastolic depolarization (Hof et al. 2013, Simard et al. 2013). Indeed, currents from a nonselective cation channel with characteristics of TRPM4 currents have also been detected in murine sinoatrial nodes and human atrial cells (Demion, Bois, Launay and Guinamard 2007, Guinamard et al. 2004). Despite this growing body of evidence illustrating the importance of TRPM4 in cardiac conduction, the role of TRPM4 in human and murine cardiac physiology remains unclear. This study aimed to bridge this knowledge gap.

In the past, two main experimental approaches have been used to study TRPM4 function: TRPM4 inhibitors or *Trpm4* ^-/-^ mouse models, but both have substantial limitations. The first approach consists of using TRPM4 inhibitors such as flufenamic acid or the widely used drug 9-phenanthrol (Guinamard et al. 2014). However, the specificity of 9-phenantrol and flufenamic acid is disputed, as recent papers have shown that 9-phenanthrol also inhibits the calcium-activated potassium channel K_Ca_3.1 and the calcium-activated chloride channel TMEM 16A (Burris et al. 2015, Garland et al. 2015) and flufenamic acid the Ca^2+^ activated Cl^−^ channels (Gwanyanya et al. 2010). To circumvent this problem, we recently identified two new specific TRPM4 inhibitors in the frame of the NCCR TransCure consortium: 4-chloro-2-[2-(2-chloro-phenoxy)-acetylamino]-benzoic acid (CBA) and 4-chloro-2-(2-(naphthalene-1-yloxy) acetamido) benzoic acid (NBA) (Arullampalam et al. 2021, Ozhathil et al. 2018).

Although both CBA and NBA decrease the current mediated by human TRPM4 overexpressed in heterologous expression systems, surprisingly, only the NBA compound inhibits mouse TRPM4 currents in this system (Arullampalam, Preti, Ross-Kaschitza, Lochner, Rougier and Abriel 2021). These findings have yet to be confirmed in *in-vivo* or *ex-vivo* experiments. The second approach consists of *Trpm4* ^-/-^ mouse models. Two groups have already developed and characterized such models, but the findings of these two studies differed significantly (Demion, Thireau, Gueffier, Finan, Khoueiry, Cassan, Serafini, Aimond, Granier, Pasquie, Launay and Richard 2014, Mathar et al. 2010). On the one hand, Mathar *et al*. noted no functional change between male *Trpm4* ^-/-^ and WT mice under basal conditions (cardiac output, stroke work, preload, and ECG parameters) (Mathar, Vennekens, Meissner, Kees, Van der Mieren, Camacho Londono, Uhl, Voets, Hummel, van den Bergh, Herijgers, Nilius, Flockerzi, Schweda and Freichel 2010). On the other hand, Demion *et al*. found that male *Trpm4* ^-/-^ mice developed left ventricle hypertrophy, multi-level conduction blocks, increased ectopic activity, and ECG alterations (Demion, Thireau, Gueffier, Finan, Khoueiry, Cassan, Serafini, Aimond, Granier, Pasquie, Launay and Richard 2014). Although these observations are important to understand the potential roles of TRPM4 for cardiac function, they also illustrate the high level of complexity of investigating the cardiac role of TRPM4 in a physiological environment. Possible explanations for the discrepancies are the different mouse strains used (129/SvJ for Mathar *et al*. and C57Bl/6J for Demion *et al*.) and the mouse substrain as reported by Rebekka *et al*. (Medert et al. 2020).

To address these discrepancies, we generated a third *Trpm4* knockdown mouse model, slightly different than the one used by Demion *et al*., on a C57Bl6/JRj background (Demion, Thireau, Gueffier, Finan, Khoueiry, Cassan, Serafini, Aimond, Granier, Pasquie, Launay and Richard 2014). The C57Bl6/JRj strain has the advantage of being well characterized and readily available via the laboratory Jackson (https://janvier-labs.com/fiche_produit/2-c57bl-6jrj/) to perform the different backcrossing necessary to obtain the mouse line with a genetic homogenous “pure” background. With this new *Trpm4* ^-/-^ mouse model, we first investigated whether we could confirm the previous observations done by the other groups on the role of TRPM4 in cardiac physiology (Ge et al. 2019, Siersbaek et al. 2020). Second, we assessed the influence of the age and sex of the animals on TRPM4 function. Lastly, we tested in a mouse model for the first time NBA, the new promising human and mouse TRPM4 inhibitor.

## METHODS

### *Trpm4* ^-/-^ C57BL/6JRj mouse model

A new *Trpm4* ^-/-^ mouse model has been generated for this research project. The *Trpm4* ^-/-^ (B6.Cg-Trpm4tm1.2-PG) mouse model is a global knockdown of the *Trpm4* gene by excision of exons 10 and were backcrossed on a C57Bl6/JRj background (Janvier) (Supplementary figure 1). To fulfill the “3R” criteria (*Reduce, Reuse, Refine)*, male and female *Trpm4* ^-/-^ mice and wild-type animals at different matched ages were used for all experiments. These included young (12 weeks old) and adult animals (more than 24 weeks old). Mice were housed in a pathogen-free, controlled environment (21 ± 1°C; humidity 60%; lights on 08:00 AM - 08:00 PM; food and water available *ad libitum*; enriched environment) with maximum 5 mice per cage. According to the Swiss Federal Animal Protection Law, all animal experiments were performed and approved by Bern’s Cantonal Veterinary Administration. This investigation conforms to the Guideline for the Care and Use of Laboratory Animals, published by the USNational Institutes of Health (NIH publication no. 85–23, revised 1996).

### Western blot

The expression of the TRPM4 channel was assessed in whole-cell lysates. First, heart tissue was lysed for 1 hour at 4 °C in lysis buffer (50 mM HEPES pH 7.4, 1.5 mM MgCl_2_, 150 mM NaCl, 1 mM EGTA pH 8, 10% glycerol, 1% Triton X-100, and Complete® protease inhibitor cocktail (Roche Diagnostics, Mannheim, Germany)). Then it was centrifuged at 4 °C, 16’000 g for 15 minutes, and centrifuged the pellet was discarded. The protein concentration of each of the lysate samples was measured in triplicate by Bradford assay and interpolated by a bovine serum albumin (BSA) standard curve. Samples were denatured at 95 °C for 5 minutes before loading them on a gel. Sixty µg of protein for each sample was run at 150 V for 1 hour on 9% polyacrylamide gels. The Turbo Blot dry blot system (Biorad, Hercules, CA, USA) was used to transfer the samples to a nitrocellulose membrane. All membranes were stained with Ponceau as a qualitative check for equivalent loading of total protein. Membranes were then rinsed twice with PBS before using the SNAP i.d. system (Millipore, Zug, Switzerland) for western blotting. The membrane was blocked with 0.1% BSA in PBS for 10 minutes. The membranes were incubated for 10 minutes with rabbit primary anti-mouse Trpm4 (Q7TN37-1) antibody (epitope: _2_VGPEKEQSWIPKIFRKKVC_10_) (generated by Pineda, Berlin, Germany) diluted 1:750 in PBS + 0.1% Tween using the SNAP i.d. system (Millipore, Billerica, MA, USA). Membranes were subsequently washed 4 times in PBS + 0.1% Tween before incubating with fluorescent secondary antibodies. Secondary antibodies (IR Dye 800 CW, 1:1000 in PBS/Tween, LI-COR Biosciences, Lincoln, NE, USA) were added for 10 minutes. After 4 more washes with PBS + 0.1% Tween and 3 washes in PBS, membranes were scanned with the Odyssey® Infrared Imaging System (LI-COR Biosciences, Bad Homberg, Germany) for detection of fluorescent protein. Subsequent quantitative analysis of protein content was achieved by measuring and comparing band densities (equivalent to fluorescence intensities of the bands) using Odyssey software version 3.0.21.

### Surface Electrocardiograms

Three-lead surface electrocardiograms (ECG) were recorded during 1 minute at 5-, 6-, 7-, 8-, 9-, 11-, 12-, and above 24-weeks-old mice under anesthesia (isoflurane IsofloH, ABBOTT S.A, Madrid, Spain: induction: 2.0% in 1000 cm^3^ O_2_/minute and maintenance: 1.5 vol.% in 500 cm^3^ O_2_/minute). Body temperature was maintained at 37°C using a thermic pad. Data was collected and transmitted to a computer via an analog-digital converter (National Instruments, Austin, TX, USA). Recordings were made at 1500 Hz. Data from lead II configuration were analyzed offline byLabChart7 Pro (AD Instruments, Castle Hill, NSW, Australia). Firstly, the ECG tracing was scanned for arrhythmias and noise. Heart rate (HR), P duration, RR, PR, and QRS intervals were determined by analyzing three stable sequences of 30 seconds for each ECG Heart rate variability was quantified using the root mean square of successive differences between normal heartbeats (RMSSD). RMSSD is obtained by first calculating each successive time difference between heartbeats. Then, each of the values is squared and the result is averaged before the square root of the total is obtained.

### Echocardiography

Each anesthetized mouse (isoflurane IsofloH, ABBOTT S.A, Madrid, Spain; induction: 3.5% in 1000 cm^3^ O_2_/minute; maintenance: 1-1.5% in 1000 cm^3^ O_2_/minute) was placed in supine position on a heated table and secured to the table via 4 ECG leads attached to the mouse’s limbs. Hair-removing cream (Veet, Reckitt Benckiser, Granollers, Spain) was used to remove chest hair, and ultrasound gel (Quick Eco-Gel, Lessa, Barcelona, Spain) was applied to enhance image quality. Echocardiography studies were done at baseline, using a Vevo 2100 ultrasound system (Visual Sonics, Toronto, Canada) equipped with a real-time micro-visualization scan head probe (MS-550D) at around 740 frames per second. The HR and respiratory rates were monitored during the study. To assess the function of the heart, left ventricle (LV) systolic, LV diastolic, and right ventricle (RV) systolic function were assessed as described below. LV systolic function was assessed via the stroke volume, the ejection fraction, the cardiac output, the left ventricle mass corrected, and the shortening fraction parameters calculated from M-mode measurement. The M-mode cursor was positioned vertically to obtain a trans-thoracic parasternal short-axis view. This classical LV M-mode tracing approach allowed the visualization of both papillary muscles. The functional parameters of the heart were calculated using the following equations based on LV diameter measurements (adapted from American Society of Echocardiography guidelines and the Vevo 2100 protocol-based measurements and calculations guide):

#### Left ventricular end-diastolic volume (LVEDV) (µL)

LVEDV=(7/(2.4+LVEDD))*LVEDD^3^

#### Left ventricular end-systolic volume (LVESV) (µL)

LVESV=(7/(2.4+LVESD))*LVESD^3^

#### Stroke volume (SV) (µL)

SV=LVEDV-LVESV

#### Cardiac output (CO) (mL/minute)

CO=SV*HR

HR: heart rate (beat per minute (bpm)).

#### Fractional shortening (%)

FS=((LVEDD–LVESD)*100)/LVEDD

LVEDD: left ventricle internal dimensions at end diastole (cm).

LVESD: left ventricle internal dimensions at end systole (cm).

#### Left ventricular ejection fraction (LVEF) (%)

LVEF=((LVEDV-LVESV)*100)/LVEDV

LVEDV: left ventricle end-diastolic volume (mL).

LVESV: left ventricle end-systolic volume (mL).

#### Left ventricle mass corrected (LV mass corrected) (mg)

LV mass corrected=((1.053*((LVEDD+LVPWED+IVSED)^3^-LVEDD^3^))*0.8)

LVPW, ED: left ventricle posterior wall of the at the end diastole (cm). IVS, ED: interventricular septum thickness at the end systole (cm) thickness of the interventricular septum. For the left ventricle mass corrected equation, the 1.053 g/mL value corresponds to the estimated density of the ventricle (Vinnakota and Bassingthwaighte 2004). The factor 0.8 is a “correction” factor applied based on initial recommendations by the American Society of Echocardiography due to the overestimation in echocardiography measurements. The left ventricle diastolic function was assessed using pulse-wave Doppler imaging of trans-mitral inflow. This was measured from the LV apical four-chamber views. The transducer was angled so that the ultrasound waves would be parallel to the blood flow. Velocities of peak E wave (early filling wave) and peak A wave (late atrial contraction wave), mitral valve E/A ratio (MV E/A), deceleration time (MV decel), left ventricle isovolumetric relaxation time (IVRT), and mitral valve pressure half time (MV PHT) were calculated based on the Doppler graph.

#### Mitral valve E/A ratio (MV E/A)

MV E/A: ratio between velocities of peak E wave (early filling wave) and peak A wave (late atrial contraction wave).

#### Mitral valve pressure half time (MV PHT) (simplified) (ms)

MV PHT=TV_max_ /1.4-TV_max_

TV_max_ corresponds to the time from the V_max_ velocity.

Mitral valve deceleration time (MV decel) (mm/s^2^):

MV deceleration time corresponds to the time point from the V_max_ to the time point where the velocity is equal to zero.

#### Left ventricle isovolumetric relaxation time (IVRT) (ms)

The isovolumic relaxation time is the time interval between the end of aortic ejection and the beginning of ventricular filling.

The right ventricle systolic function was also assessed using the pulmonary artery view via the pulse-wave Doppler imaging of trans-pulmonary valve inflow. The Doppler graph was used to measure the peak velocity of blood flow at the pulmonary valve (PV peak vel), peak pressure (PV peak pressure), diameter (PV diam), and mean pulmonary artery pressure (mPAP).

#### Pulmonary valve peak velocity (PV peak vel) (mm/s)

Correspond to the interval between the onset of flow and peak flow.

#### Pulmonary valve diameter (PV diam) (mm)

Correspond to the diameter of the pulmonary valve.

#### Pulmonary valve peak pressure (PV peak pressure) (mmHg)

Correspond to the maximum pressure at the level of the pulmonary artery.

#### Mean pulmonary artery pressure-mPAP (mmHg)

mPAP=90-(0.62*AT_RVOT_)

AT_RVOT_ corresponds to the acceleration time of the right ventricular outflow tract measured from the beginning of the flow to the peak flow velocity.

### Pseudo- and intracardiac ECG

Anesthetized mice (ketamine (200mg/kg) and xylazine (20mg/kg)) were sacrificed by cervical dislocation. Hearts were rapidly excised and retrogradely perfused using the Langendorff system (modified Krebs-Henseleit buffer (KHB) containing the following in concentrations of mmol/L:116.5 NaCl_2_, 25 NaHCO_3_, 4.7 KCl, 1.2 KH_2_PO_4_, 11.1 glucose, 1.5 CaCl2, 2 Na-pyruvate and bubbled with 95% O_2_ and 5% CO_2_ at 37°C). Perfusion pressure was maintained at 70 mmHg. Pseudo-ECGs were recorded from the outer side of the explanted heart. Negative and positive ends were placed at the root of the aorta and apex of the ventricles, respectively. Intracardiac ECGs of the “right heart” (right atrium and right ventricle) were measured from the inner side of the explanted heart using octopolar silver electrodes. The closest derivation to the atrioventricular node was used for the electrical activation of the right atrium (A wave) (“Ch5” in figure 6A) and the derivation at the apex of the right ventricle for the electrical activation of the right ventricle (V wave) (“Ch1” in figure 6A). Power lab bio-amplifiers collected the ECG tracings, whic were low- and high-pass filtered (at 5 kHz and 10 kHz, respectively) and sampled at a speed of 1 k/s. Initially, the heart was perfused with KHBfor the first 10 minutes, after which a solution of 5 µM NBA, 5 µM CBA or vehicle control (DMSO 0.05%) dissolved in KHB was added to the perfusion. The ECG tracings obtained were analyzed using Lab Chart Pro V8. (AD. Instruments, Australia) to determine the P wave, QRS complex, and QT region. Visualization of at least six intracardiac ECG leads at the level of the right atrium and right ventricle establishes the correct position of the catheter. Atrioventricular delay was measured between channels 5 and 6 as it displayed waves A and V at the Ch1.

### RNA preparation and real-time quantitative RT-PCR

Isolated RNA from atrial and ventricular myocytes was extracted and amplified using PCR techniques. Both left and right atrial myocytes were used from 3 mice for each genotype. TaqMangene expression assay probes for mouse Trpm4 (Mm01205532-m1) and GAPDH (Mm99999915-g1) were used to amplify the isolated RNA Comparative threshold (CT) values and target *Trpm4* genes were determined for reference for each sample set. Relative quantification was then performed using the CT method (ΔΔCT).

### Cell line preparations, transfection, and whole-cell electrophysiology

Human embryonic kidney (HEK-293) cells were cultured in DMEM (Gibco, Basel, Switzerland) supplemented with 10% FBS, 0.5% penicillin, and streptomycin (10,000 U/mL) at 37°C in a 5% CO_2_ incubator. For electrophysiological studies, T25 flasks with HEK-293 cells were transiently co-transfected using X-tremeGENE™ 9 reagent (Sigma-Aldrich, Darmstadt, Germany) with 2 µg of mouse Na_v_1.5 cDNA. All transfections included 0.1 µg of cDNA encoding the GFP protein. Forty-eight hours post-transfection, green cells were selected and analyzed for patch clamp experiments. Peak sodium currents (*I*_Na_) were recorded in the whole-cell configuration at RT (22-23°C) using a VE-2 amplifier (Alembic Instrument, USA). Sodium currents were digitized at a sampling frequency of 20 kHz. Borosilicate glass pipettes were pulled to a series resistance of ∼2 MΩ. pClamp software, version 8 (Axon Instruments, Union City, CA, USA) was used for recordings. Data were analyzed using pClamp software, version 8 (Axon Instruments). HEK-293 cells were bathed in a solution containing (in mM) NaCl 30, NMDG-Cl 100, CaCl_2_ 2, MgCl_2_ 1.2, CsCl 5, HEPES 10, and glucose 5 (pH was adjusted to 7.4 with CsOH). HEK-293 cells were initially voltage-clamped (holding potential -80 mV) and dialyzed with an internal solution containing (in mM) CsCl 60, Cs-aspartate 70, CaCl_2_ 1, MgCl_2_ 1, HEPES 10, EGTA 11, and Na_2_ATP 5 (pH was adjusted to 7.2 with CsOH). A pulse protocol consisting to a depolarization step for 20 ms from -80 mV to -20 mV was applied every 3 seconds. The sodium peak current was first recorded under vehicle perfusion (DMSO 0.05%) for 1 minute (or until stability was reached). Then, the compound (NBA or CBA 5 µM) or vehicle (DMSO 0.05%) was applied for 5 minutes (or until recordings had stabilized). The sodium peak currents measured during the stable phase during the vehicle, and compound/vehicle perfusion were used to quantify the percentage of the effect of each compound or the vehicle using the following formula: Percentage of effect on sodium current=100-(100\**I*_Na compounds/vehicle_/*I*_Na vehicle_).

### Drugs

Stock solutions of 4-chloro-2-(2-(naphthalene-1-yloxy) acetamido) benzoic acid (NBA) and 4-chloro-2-[2-(2-chloro-phenoxy)-acetylamino]-benzoic acid (CBA) were made in 100% DMSO to a final concentration of 10 mM. Freshly diluted NBA and CBA solutions at 5 µM (DMSO 0.05%) were made before each experiment using either Krebs-Henseleit buffer (intracardiac ECG) or extracellular solution (patch clamp). The final concentration of NBA was chosen based on the previously determined concentration estimated to fully block mouse TRPM4 channel overexpressed in a heterologous expression system (Arullampalam, Preti, Ross-Kaschitza, Lochner, Rougier and Abriel 2021). Given that CBA failed to inhibit mouse TRPM4 in a heterologous expression system, this compound was chosen as a negative control for experiments (Arullampalam, Preti, Ross-Kaschitza, Lochner, Rougier and Abriel 2021).

### Data analyses and statistics

Data are represented as means ± SEM Statistical analyses were performed using Prism7 GraphPad™ software. An unpaired nonparametric t-test followed by Mann-Whitney U post-test was used to compare two unpaired groups. A paired nonparametric t-test followed by a Wilcoxon matched-pairs signed-rank post-test was used to compare two paired groups. *p* < 0.05 was considered significant.

## RESULTS

### Expression of TRPM4 in the heart

The role of TRPM4 in cardiac function has been investigated with a new *Trpm4* ^-/-^ mouse model, in which exon 10 of the *Trpm4* gene has been deleted (Supplementary figure 1) (Ozhathil et al. 2021).). Like the two other models, the *Trpm4*^-/-^ mouse model did not show any increase in mortality or alteration of Mendelian genetic transmission. At a young age, body weight averages for both sexes were similar. However, at adult age, body weight was 9.0% lower only in adult male *Trpm4* ^-/-^ compared to their controls ((average body weight _control adult male_: 33.21 ± 0.62 gram (n = 28) and average body weight _*Trpm4* -/- adult male_: 30.21 ± 0.60 gram (n = 26); p<0.01) and (average body weight _control adult female_: 25.43 ± 0.99 gram (n = 25) and average body weight _*Trpm4* -/- adult female_: 24.03 ± 0.45 gram (n = 24); p>0.05)). As expected, cardiac TRPM4 expression in *Trpm4* ^-/-^ hearts is drastically decreased in the heart compared to controls, both at the mRNA level (Figure 1A) and at the protein level (Figures 1B and 1C) as previously reported (Ozhathil, Rougier, Arullampalam, Essers, Ross-Kaschitza and Abriel 2021).

**Figure 1.**
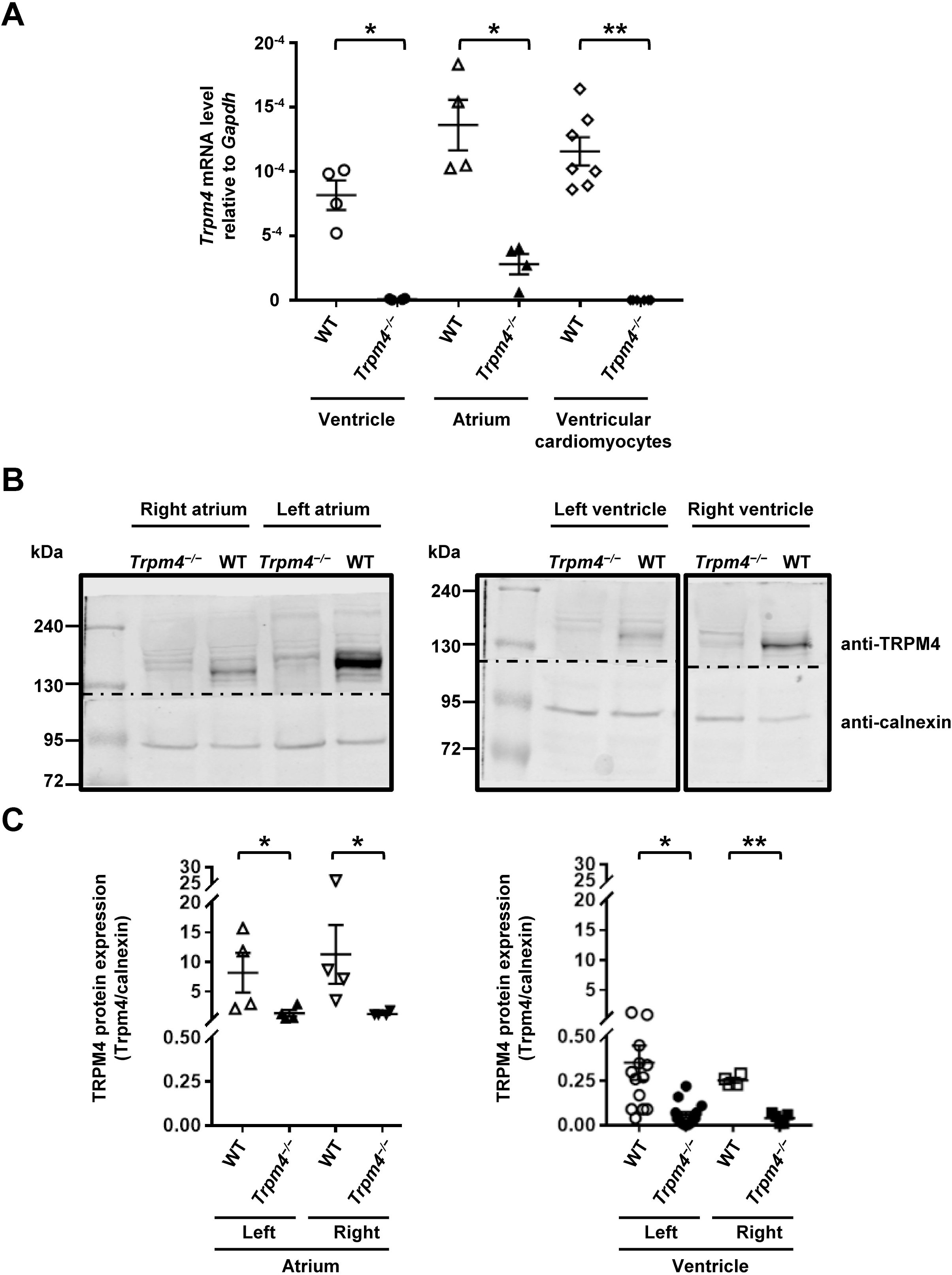
Trpm4 expression in mouse heart. *A*: Quantification of *Trpm4* mRNA using RT-qPCR. For the ventricle, atrium, and ventricular cardiomyocytes, Ct values were corrected with the housekeeping gene *Gapdh*. *: p⩽0.05 and **: p⩽0.01 (n⩾4 per group). *B*: Expression of Trpm4 in WT and *Trpm4* ^-/-^ mouse atrial and ventricular tissues. Representative immunoblots of Trpm4 protein expression from the right atrium, left atrium, right ventricle, and left ventricle. Dotted lines indicate cut in western blot membranes. *C*: Quantification of the immunoblots showing protein expression from the right atrium, left atrium, right ventricle, and leftventricle (Y-axis is split for clarity). *: p⩽0.05 and **: p⩽0.01 (n⩾4 per group).

### Assessment of electrophysiological parameters by classic ECG

Surface ECGs were recorded to assess the cardiac electrophysiology of eight groups of mice (young WT male and female, young *Trpm4* ^-/-^ male and female, adult WT, and female, and adult *Trpm4* ^-/-^ male and female) (Figure 2A). Before these ECGs were performed, cardiac electrical parameters were first characterized in male and female *Trpm4* ^-/-^ and WT mice by recording weekly ECGs in 5- to 9-weeks-old animals (Table 1). These ECGs did not reveal any differences in the measured parameters (RR interval, PR interval, QRS interval, and P duration) between the different groups (Table 1). Moreover, most of the investigated ECG parameters did not differ when comparing young and adult mice of both sexes and with wild-type and *Trpm4* ^-/-^ genetic backgrounds (Table 1). Only a significant heart rate decreases of 8 % ± 2 % (n = 21), with a concomitant increase of the RR interval, was observed in male adult *Trpm4* ^-/-^ mice compared to male adult WT (Figure 2B, and Table 1). Interestingly, quantifying the heart rate variability (HRV) using the RMSSD index showed that the knockdown of the *Trpm4* gene increases HRV only in adult male *Trpm4* ^-/-^ hearts (Figure 2C and Table 1).

**Figure 2.**
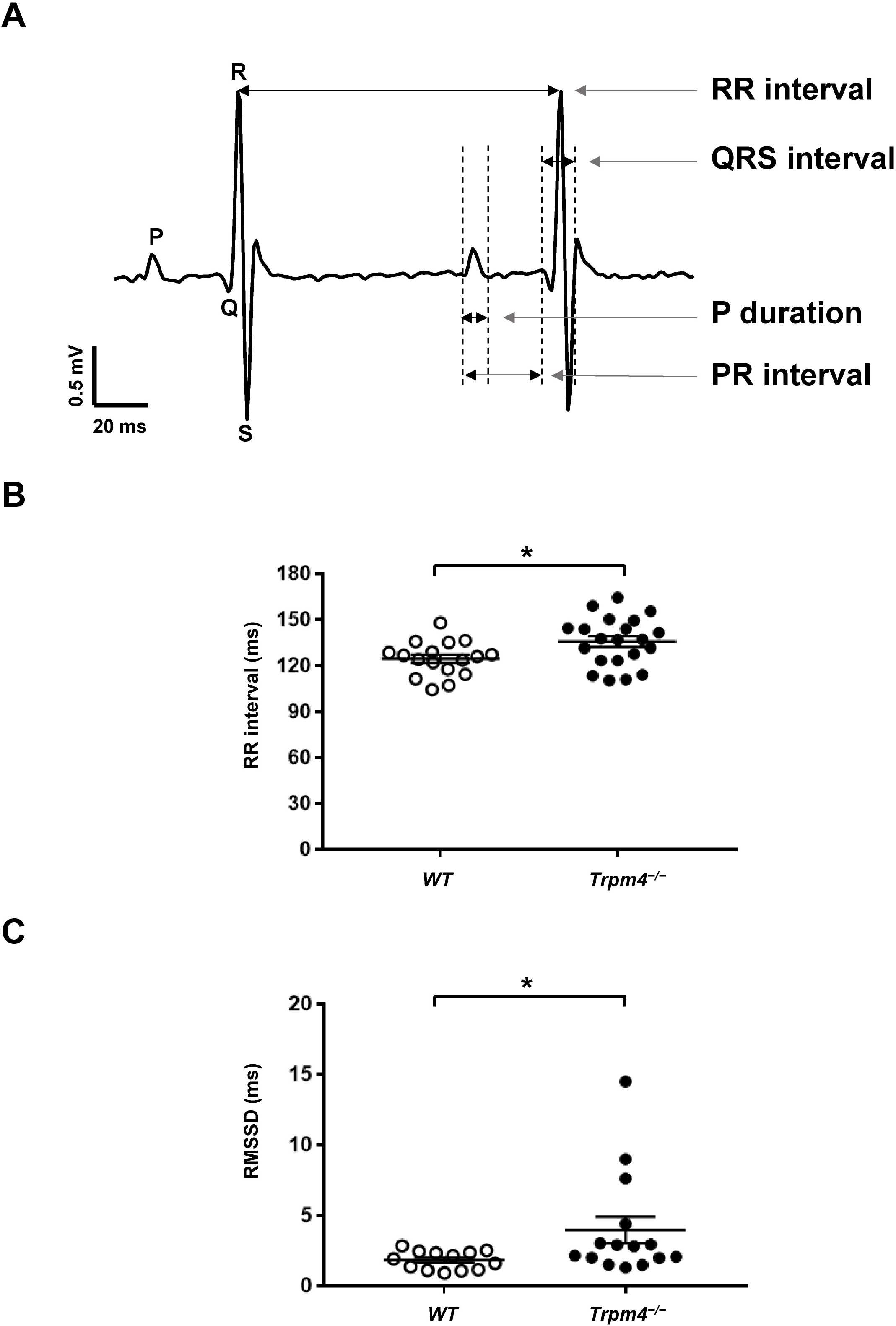
ECG parameters in WT and *Trpm4* ^-/-^ mice. *A*: Representative E.C.G. trace from adult male wild-type mouse and the different parameters investigated. *B* and *C*: RR interval duration (*B*) and RMSSD parameter (*C*) showing the difference between 24-week-old male WT and*Trpm4* ^-/-^ mice. n.d: not determined, and *: p⩽0.05 (n⩾13 per group).

**Table 1.** Evolution of the ECG parameters at 5-, 6-, 7-, 8-, 9-weeks-old of age, at a young age (12-weeks-old) and adult age (more than 24-weeks-old). ECG parameters from male and female WT and *Trpm4* ^*-/-*^ mice at different ages. Values highlighted in light grey (*: p⩽0.05) indicate a statistical difference between *Trpm4* ^-/-^ and control for the respective matching age and sex group.

|  |  | RR interval (ms) |  |  |  |  |  |  |
| --- | --- | --- | --- | --- | --- | --- | --- | --- |
|  |  | 5<sup>th</sup> week | 6<sup>th</sup> week | 7<sup>th</sup> week | 8<sup>th</sup> week | 9<sup>th</sup> week | Young | Adult |
| control | ♂ | 136.5±3.0<br>(n=4) | 119.2±4.8<br>(n=4) | 128.7±2.5<br>(n=4) | 124.5±0.9<br>(n=4) | 124.6±2.2<br>(n=4) | 122.6±2.4<br>(n=16) | 124.5±2.7<br>(n=17) |
| *Trpm4*<sup>-/-</sup> | ♂ | 135.8±3.8<br>(n=5) | 127.6±5.1<br>(n=5) | 117.7±3.5<br>(n=5) | 127.6±5.4<br>(n=5) | 120.4±2.9<br>(n=5) | 131.6±3.6<br>(n=13) | 135.7±3.5<br>(n=21) |
| control | ♀ | 144.8±3.2<br>(n=4) | 121.5±5.5<br>(n=4) | 124.2±2.7<br>(n=4) | 117.9±4.8<br>(n=4) | 107.3±1.8<br>(n=4) | 127.1±2.8<br>(n=11) | 137.6±3.2<br>(n=7) |
| *Trpm4*<sup>-/-</sup> | ♀ | 151.3±12.6<br>(n=3) | 145.5±2.4<br>(n=3) | 133.8±12.0<br>(n=3) | 131.5±2.0<br>(n=3) | 134.1±5.6<br>(n=3) | 133.4±7.7<br>(n=8) | 143.8±3.5<br>(n=9) |
|  |  | PR interval (ms) |  |  |  |  |  |  |
|  |  | 5<sup>th</sup> week | 6<sup>th</sup> week | 7<sup>th</sup> week | 8<sup>th</sup> week | 9<sup>th</sup> week | Young | Adult |
| control | ♂ | 34.8±1.0<br>(n=4) | 31.7±1.7<br>(n=4) | 35.0±1.8<br>(n=4) | 35.8±1.5<br>(n=4) | 35.9±1.4<br>(n=4) | 39.5±0.6<br>(n=16) | 40.5±0.7<br>(n=17) |
| *Trpm4*<sup>-/-</sup> | ♂ | 37.6±1.2<br>(n=5) | 38.1±0.6<br>(n=5) | 37.2±0.8<br>(n=5) | 38.0±0.7<br>(n=5) | 36.6±0.8<br>(n=5) | 40.4±0.7<br>(n=13) | 41.2±0.7<br>(n=21) |
| control | ♀ | 37.5±1.1<br>(n=4) | 36.5±0.6<br>(n=4) | 36.6±0.6<br>(n=4) | 37.8±2.7<br>(n=4) | 39.6±1.6<br>(n=4) | 40.3±1.2<br>(n=11) | 41.8±1.0<br>(n=7) |
| *Trpm4*<sup>-/-</sup> | ♀ | 35.9±1.8<br>(n=3) | 38.1±1.3<br>(n=3) | 36.2±0.8<br>(n=3) | 38.6±1.7<br>(n=3) | 35.9±0.4<br>(n=3) | 40.2±1.7<br>(n=8) | 42.3±1.0<br>(n=9) |
|  |  | QRS interval (ms) |  |  |  |  |  |  |
|  |  | 5<sup>th</sup> week | 6<sup>th</sup> week | 7<sup>th</sup> week | 8<sup>th</sup> week | 9<sup>th</sup> week | young | Adult |
| control | ♂ | 10.5±0.8<br>(n=4) | 9.8±0.5<br>(n=4) | 11.2±0.5<br>(n=4) | 10.7±0.5<br>(n=4) | 11.2±0.0<br>(n=4) | 10.4±0.4<br>(n=16) | 10.1±0.3<br>(n=17) |
| *Trpm4*<sup>-/-</sup> | ♂ | 10.5±0.5<br>(n=5) | 10.3±0.6<br>(n=5) | 10.4±0.6<br>(n=5) | 10.5±0.5<br>(n=5) | 11.1±0.3<br>(n=5) | 9.8±0.4<br>(n=13) | 10.9±0.3<br>(n=21) |
| control | ♀ | 10.7±0.6<br>(n=4) | 10.6±0.4<br>(n=4) | 10.9±0.2<br>(n=4) | 10.4±0.5<br>(n=4) | 10.8±0.5<br>(n=4) | 10.9±0.3<br>(n=11) | 11.0±0.5<br>(n=7) |
| *Trpm4*<sup>-/-</sup> | ♀ | 11.8±0.6<br>(n=3) | 11.2±0.8<br>(n=3) | 11.3±0.1<br>(n=3) | 12.3±0.3<br>(n=3) | 11.4±0.5<br>(n=3) | 11.2±0.5<br>(n=8) | 10.2±0.5<br>(n=9) |
|  |  | P duration (ms) |  |  |  |  |  |  |
|  |  | 5<sup>th</sup> week | 6<sup>th</sup> week | 7<sup>th</sup> week | 8<sup>th</sup> week | 9<sup>th</sup> week | Young | Adult |
| control | ♂ | 8.8±0.5<br>(n=4) | 8.4±0.4<br>(n=4) | 10.4±0.4<br>(n=4) | 9.8±0.3<br>(n=4) | 10.0±0.5<br>(n=4) | 16.7±1.2<br>(n=16) | 14.4±1.0<br>(n=17) |
| *Trpm4*<sup>-/-</sup> | ♂ | 11.0±0.7<br>(n=5) | 11.0±0.8<br>(n=5) | 10.4±0.4<br>(n=5) | 11.1±0.4<br>(n=5) | 10.7±0.6<br>(n=5) | 15.0±1.2<br>(n=13) | 17.1±1.3<br>(n=21) |
| control | ♀ | 10.4±0.3<br>(n=4) | 9.3±0.5<br>(n=4) | 9.8±0.5<br>(n=4) | 10.0±1.2<br>(n=4) | 8.4±0.8<br>(n=4) | 13.5±1.5<br>(n=11) | 19.0±1.0<br>(n=7) |
| *Trpm4*<sup>-/-</sup> | ♀ | 10.9±0.3<br>(n=3) | 11.1±0.9<br>(n=3) | 11.1±0.6<br>(n=3) | 11.9±0.6<br>(n=3) | 10.3±0.7<br>(n=3) | 16.3±2.0<br>(n=8) | 15.3±1.7<br>(n=9) |
|  |  | RMSSD (ms) |  |  |  |  |  |  |
|  |  | 5<sup>th</sup> week | 6<sup>th</sup> week | 7<sup>th</sup> week | 8<sup>th</sup> week | 9<sup>th</sup> week | Young | Adult |
| control | ♂ | n.d | n.d | n.d | n.d | n.d | 6.5±4.2<br>(N=7) | 1.8±0.2<br>(N=13) |
| *Trpm4*<sup>-/-</sup> | ♂ | n.d | n.d | n.d | n.d | n.d | 2.3±0.3<br>(N=10) | 4.0±0.9<br>(N=15) |
| control | ♀ | n.d | n.d | n.d | n.d | n.d | 6.6±2.8<br>(N=10) | 7.6±3.2<br>(N=6) |
| *Trpm4*<sup>-/-</sup> | ♀ | n.d | n.d | n.d | n.d | n.d | 3.2±1.3<br>(N=7) | 3.0±0.7<br>(N=8) |

### Characterization of left ventricular structure and function by echocardiography

To characterize the left ventricle, we studied diastolic and systolic functions (stroke volume, ejection fraction, fractional shortening, cardiac output, isovolumetric conduction, mitral valve pressure half time simplified, and left ventricle mass corrected) of *Trpm4* ^-/-^ and control mice using echocardiography. The systolic function of the left ventricle was evaluated using the parasternal short-axis view of the M mode (Figure 3A). The diastolic function was assessed using power Doppler imaging to obtain an apical four-chamber view (Figures 4A, and 5A). Scans of the hearts of young animals revealed that the mitral valve pressure half-time was significantly increased in *Trpm4* ^-/-^ male animals compared to their controls (Table 2). Other parameters did not differ between control and *Trpm4* ^-/-^ animals for both sexes (Table 2). At adult age, however, *Trpm4* ^-/-^ males presented with a higher left ventricular mass (corrected values) than controls, which correlated with slightly lower cardiac output than their respective controls (Figures 3B, 3C, and Table 2). The ejection fraction, which reflects the systolic function, was unaltered in male adult *Trpm4* ^-/-^ animals compared to their controls (Figure 3D and Table 2). Conversely, the left ventricle mass corrected values of *Trpm4* ^-/-^ females tend to be smaller than their controls (Table 2). These results suggest that male adult *Trpm4* ^-/-^ animals tend to develop left ventricle hypertrophy with age without systolic dysfunction.

**Figure 3.**
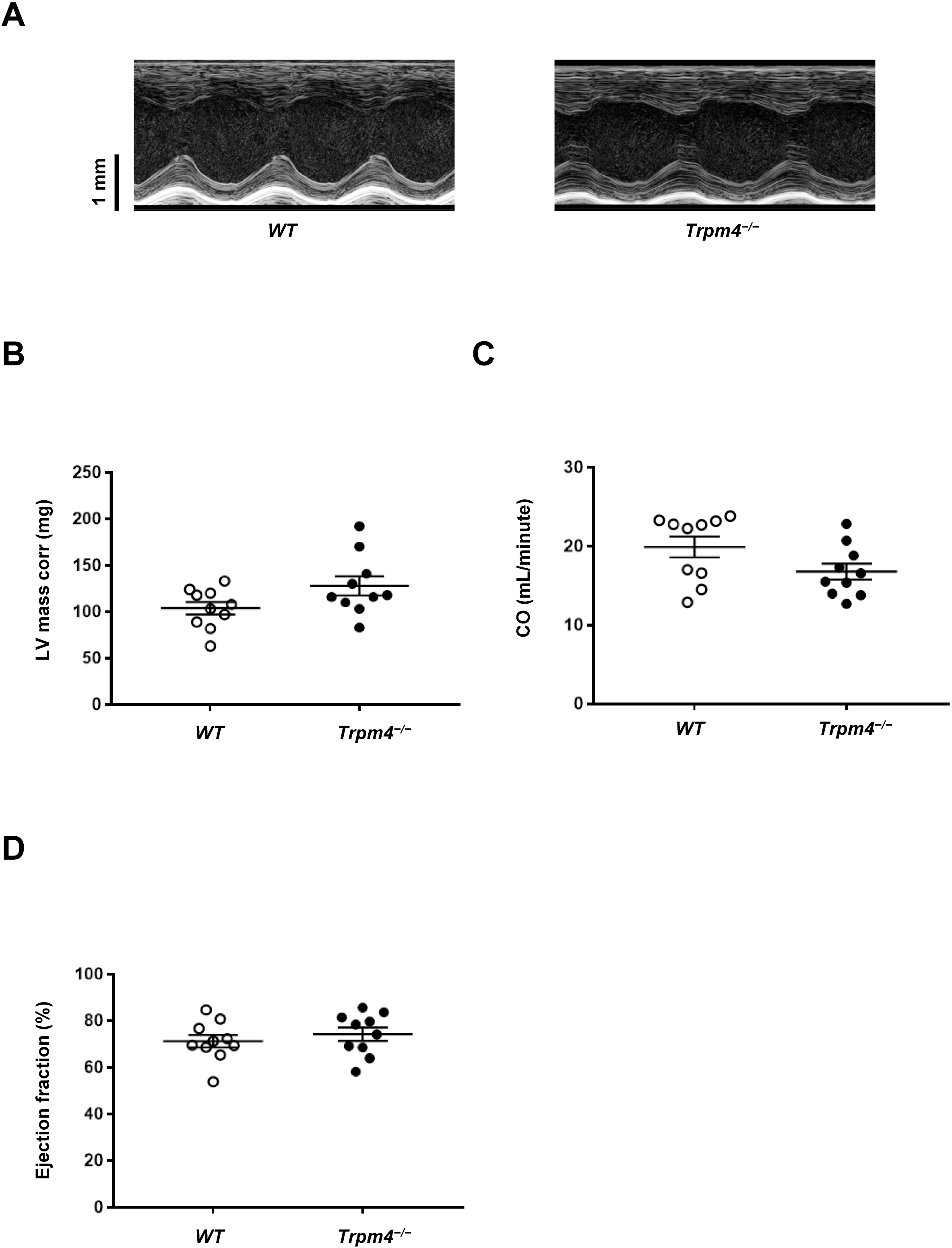
Left ventricle structure and function of WT and *Trpm4* ^-/-^ mouse hearts. *A*: Raw captions from WT and *Trpm4* ^-/-^ mouse hearts from the parasternal, short axis, M-mode echocardiography. *B, C*, and *D*: left ventricle mass corrected (LV Mass corr), cardiac output (CO), and ejection fraction parameters recorded in adult male WT and *Trpm4* ^-/-^ mice suggesting an increase of the left ventricle mass of *Trpm4* ^-/-^ mice (*B*) and a decrease of the cardiac output (*C*) without systolic dysfunction (*D*) (n=10 per group).

**Figure 4.**
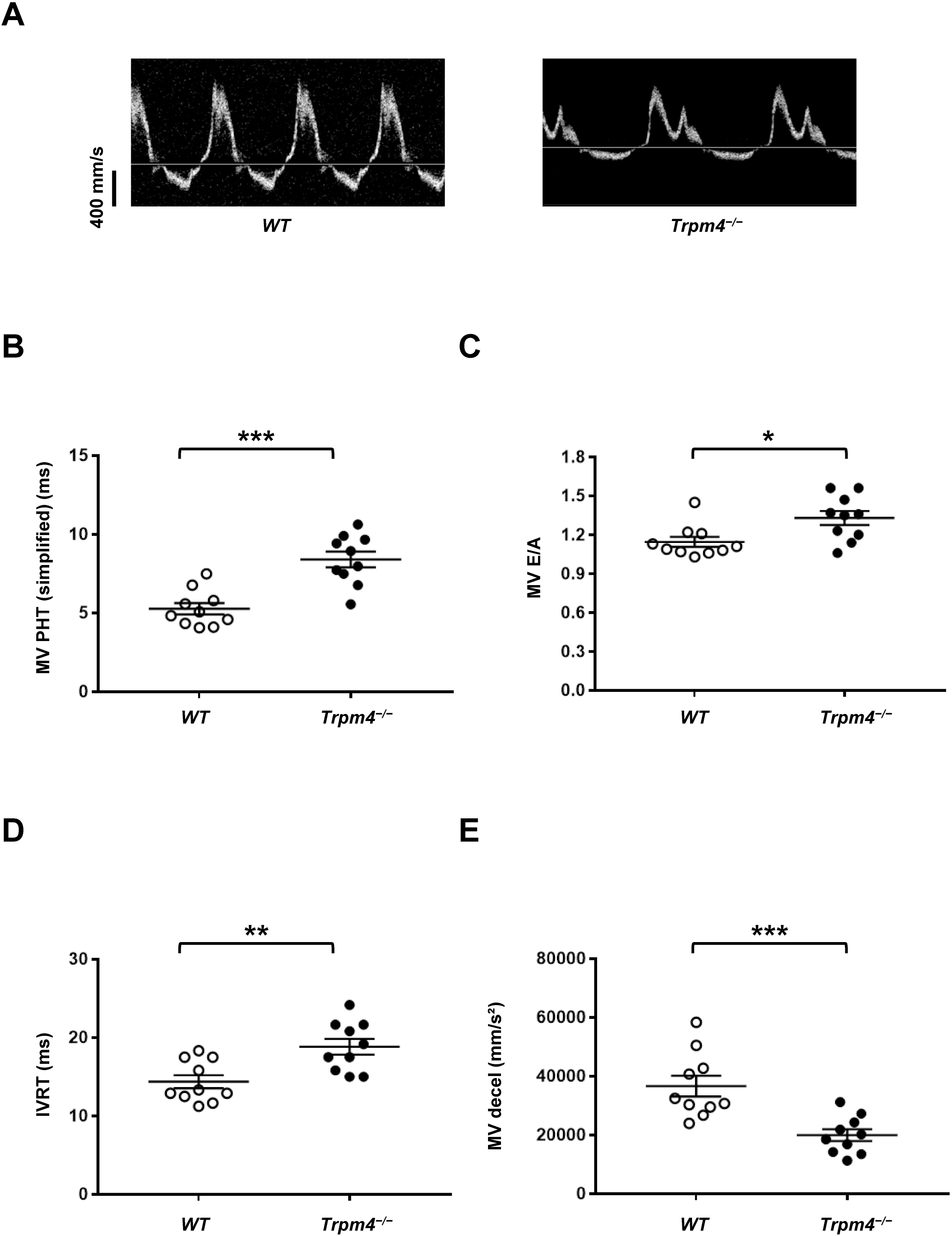
Analysis of mitral valve function of WT and *Trpm4* ^-/-^ mouse hearts. *A*: Traces representing the blood flow velocity through the mitral valves of male WT (left) and *Trpm4* ^-/-^ (right) hearts. *B, C, D*, and *E*: Dot plots showing the mitral valve pressure half time (MV PHT) (*B*), the mitral valve E/A ratio (MV E/A), which correspond to the ratio between the velocities of E wave (early filling) and A wave (atrial filling) of the mitral valve (*C*), the mitral valve isovolumetric relaxation time (IVRT) (*D*), and the mitral valve deceleration time MV (decel) (*E*). *: p⩽0.05, **: p⩽0.01, and ***: p⩽0.001 (n=10 per group).

**Figure 5.**
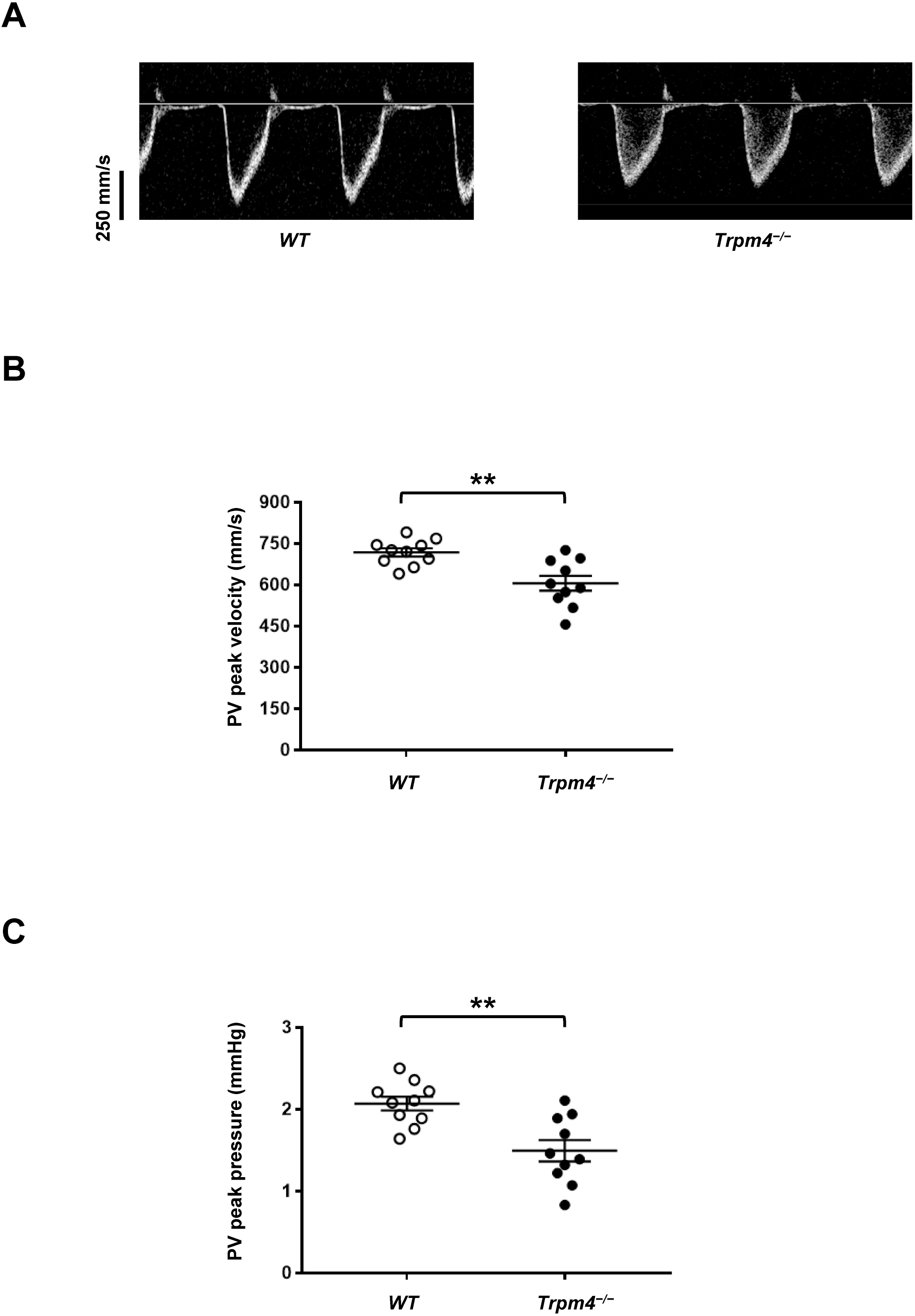
Analysis of pulmonary artery blood flow of WT and *Trpm4* ^-/-^ mouse hearts. *A*: Traces representing the velocity of blood flow in the pulmonary artery of WT (left) and *Trpm4*^-/-^ (right) hearts. *B* and *C*: dot blots showing the pulmonary valve peak velocity (*B*) and the pulmonary valve peak pressure (*C*). **: p⩽0.01 (n=10 per group).

**Figure 6.**
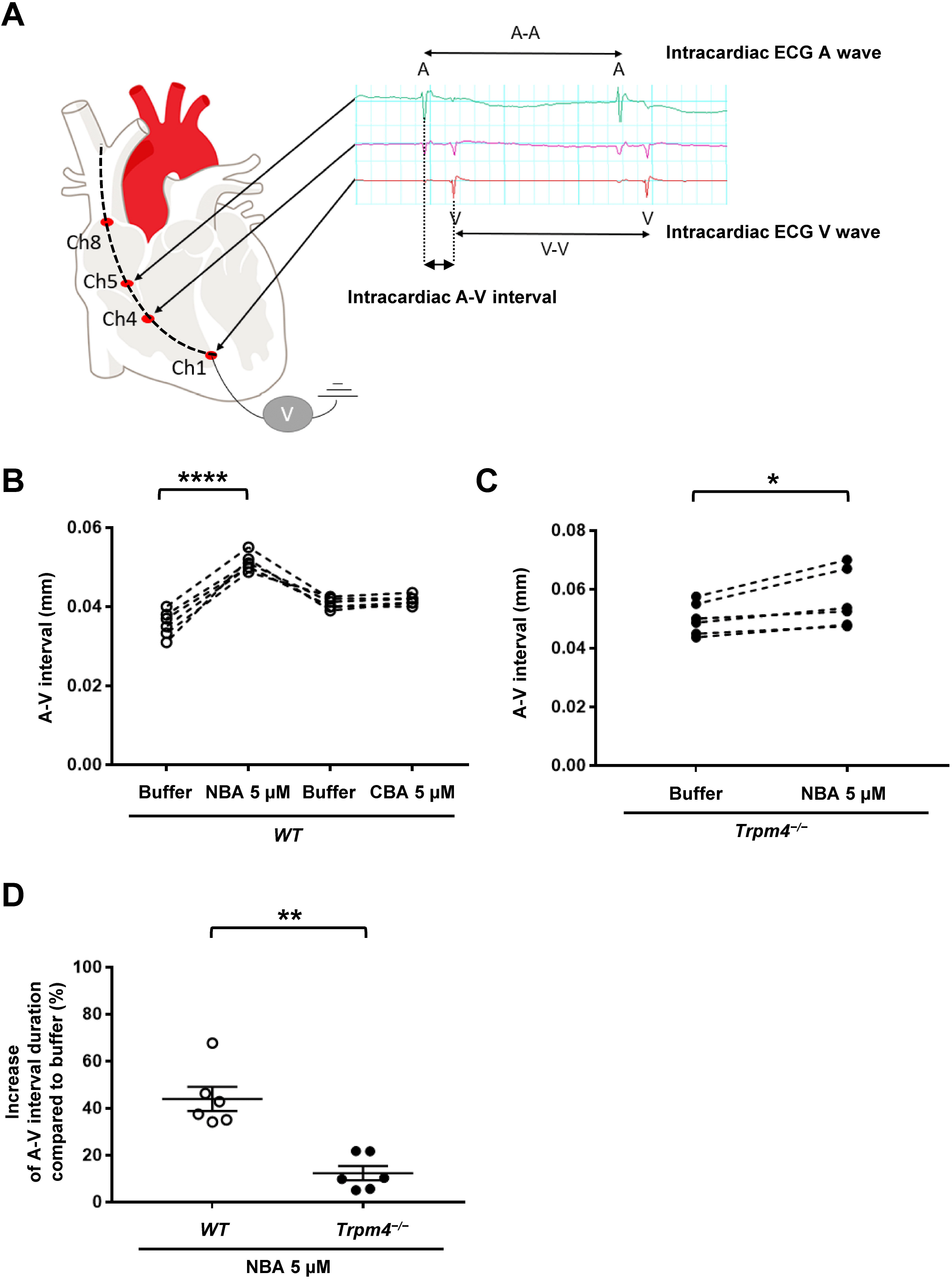
Effects of NBA and CBA on cardiac electrical activity in young male hearts using intracardiac recordings. *A*: Schema showing the localization of the octopolar probe in the “right heart” to record the A and V waves and intracardiac ECG using male wild-type heart. The ECG recording shows the A wave (right atrial electrical activity) and V wave (right ventricle electrical activity). *B* and *C*: Dot blots showing the evolution of the A-V interval after the application of 5 µM NBA on wild-type mouse hearts (*B*), 5 µM CBA on wild-type mouse hearts (*B*), and 5 µM NBA on *Trpm4* ^-/-^ mouse hearts (*C*). *D*: Dot plot summarizing the percentage of A-V interval increase between wild-type and *Trpm4* ^-/-^ mouse hearts after perfusion of 5 µM NBA *: p⩽0.05, **: p⩽0.01, and ****: p⩽0.0001 (n⩾4 per group).

**Table 2.** Echocardiography parameters related to the left ventricular structure and function. Left ventricle parameters from young and adult, male and female, and WT and *Trpm4* ^*-/-*^ mice.

|  |  | Stroke Volume (μl) | Ejection Fraction (%) | Fractional Shortening (%) | Cardiac Output (mL/minute) | LV Mass Corrected (mg) |
| --- | --- | --- | --- | --- | --- | --- |
| Young |  |  |  |  |  |  |
| control | ♂ | 39.4±5.5 (n=5) | 71.8±2.1 (n=5) | 40.3±1.8 (n=5) | 20.601±2.934 (n=5) | 94±8 (n=5) |
| *Trpm4*<sup>-/-</sup> | ♂ | 36.8±1.4 (n=5) | 72.1±4.4 (n=5) | 41.1±3.7 (n=5) | 16.297±0.821 (n=5) | 86±6 (n=5) |
| control | ♀ | 35.7±3.2 (n=4) | 74.4±2.1 (n=4) | 42.4±1.8 (n=4) | 14.647±1.338 (n=4) | 66±2 (n=4) |
| *Trpm4*<sup>-/-</sup> | ♀ | 38.8±2.9 (n=4) | 76.5±3.9 (n=4) | 44.8±3.8 (n=4) | 16.500±0.725 (n=4) | 87±13 (n=4) |
| Adult |  |  |  |  |  |  |
| control | ♂ | 43.4±1.6 (n=10) | 71.3±2.7 (n=10) | 40.4±2.2 (n=10) | 19.909±1.320 (n=10) | 104±7 (n=10) |
| *Trpm4*<sup>-/-</sup> | ♂ | 39.3±1.9 (n=10) | 74.3±2.9 (n=10) | 43.0±2.4 (n=10) | 16.754±1.021 (n=10) | 128±10 (n=10) |
| control | ♀ | 41.3±2.8 (n=8) | 69.6±2.0 (n=8) | 38.7±1.6 (n=8) | 17.930±1.185 (n=8) | 93±7 (n=8) |
| *Trpm4*<sup>-/-</sup> | ♀ | 37.3±3.1 (n=10) | 73.3±2.1 (n=10) | 41.7±1.8 (n=10) | 15.187±1.396 (n=10) | 73±8 (n=10) |

Interestingly, the remaining average mitral valve pressure (half-time simplified) was increased in male adult *Trpm4* ^-/-^ mice compared to the control animals (Figure 4B, and Table 3), suggesting a mitral valve dysfunction. Therefore, other mitral valve function parameters were recorded (mitral valve deceleration, mitral valve E/A ratio, and isovolumetric relaxation time), which were all significantly altered in adult male *Trpm4* ^-/-^ animals compared to control (Figures 4C, 4D, 4E, and Table 3). Moreover, female adult *Trpm4* ^-/-^ mice, compared to their controls, also showed alteration of different parameters related to the mitral valve function (Table 3).

**Table 3.** Echocardiography parameters related to the mitral valve structure and function. Mitral valve parameters from young and adult, male and female, and WT and *Trpm4* ^*-/-*^ mice. Values highlighted in light grey (*: p⩽0.05), dark gray (**: p⩽0.01), and black (***: p⩽0.001) indicate a statistical difference between *Trpm4* ^-/-^ and control for the respective matching age and sex group.

|  |  | MV PHT (simplified) (ms) | MV decel (mm/s<sup>2</sup>) | IVRT (ms) | MV E/A |
| --- | --- | --- | --- | --- | --- |
| Young |  |  |  |  |  |
| control | ♂ | 4.42±0.41 (n=5) | -47653±8455 (n=5) | 13.50±1.16 (n=5) | 1.30±0.06 (n=5) |
| *Trpm4*<sup>-/-</sup> | ♂ | 7.26±1.07 (n=5) | -30357±6829 (n=5) | 18.50±1.61 (n=5) | 1.38±0.03 (n=5) |
| control | ♀ | 5.80±1.11 (n=4) | -34605±5724 (n=4) | 18.06±1.47 (n=4) | 1.15±0.10 (n=4) |
| *Trpm4*<sup>-/-</sup> | ♀ | 5.31±0.61 (n=4) | -38129±6466 (n=4) | 17.50±0.90 (n=4) | 1.15±0.12 (n=4) |
| Adult |  |  |  |  |  |
| control | ♂ | 5.27±0.36 (n=10) | -36626±3526 (n=10) | 14.38±0.84 (n=10) | 1.15±0.04 (n=10) |
| *Trpm4*<sup>-/-</sup> | ♂ | 8.41±0.50 (n=10) | -19953±2010 (n=10) | 18.83±1.00 (n=10) | 1.33±0.05 (n=10) |
| control | ♀ | 4.53±0.26 (n=8) | -36026±2027 (n=8) | 14.20±1.38 (n=8) | 1.16±0.09 (n=8) |
| *Trpm4*<sup>-/-</sup> | ♀ | 7.16±0.71 (n=10) | -28216±5523 (n=10) | 18.38±1.29 (n=10) | 1.45±0.09 (n=10) |

Overall, these data suggest that, compared to their controls, male *Trpm4* ^-/-^ animals tend to develop with age left ventricle hypertrophy along with a mitral dysfunction, while female *Trpm4* ^-/-^ mice do not. Female *Trpm4* ^-/-^ mice however present similar mitral valve dysfunctions as male *Trpm4* ^-/-^ mice do.

### Characterization of the right ventricle by echocardiography

Visualizing the right ventricle with power Doppler imaging proved technically challenging as the sternum obstructed the view; therefore, we could not obtain an apical four-chamber view (Figure 5A). Instead, we indirectly investigated the right ventricle systolic function by scanning the pulmonary artery and valve, determining pulmonary valve peak velocity, pulmonary valve diameter, pulmonary valve peak pressure, and the mean pulmonary internal pressure (mPAP common). In young animals of both sexes, no alterations in the different parameters were observed (Table 4). In male adult animals, only the pulmonary valve peak velocity and the pulmonary valve peak pressure were significantly lower in male *Trpm4* ^-/-^ animals compared to their controls suggesting an alteration of the right ventricle of adult male *Trpm4* ^-/-^ animals (Figures 5B, 5C, and Table 4). No differences between groups of female adult mice were observed.

**Table 4.** Echocardiography parameters related to the “right ventricular” structure and function. Pulmonary blood flow parameters from young and adult, male and female, and WT and *Trpm4* ^*-/-*^ mice. Values highlighted in dark gray (**: p⩽0.01) indicate a statistical difference between *Trpm4* ^-/-^ and control for the respective matching age and sex group.

|  |  | PV peak velocity (mm/s) | PV diameter (mm) | PV peak pressure (mmHg) | mPAP (common) (mmHg) |
| --- | --- | --- | --- | --- | --- |
| **Young** |  |  |  |  |  |
| control | ♂ | -708±44 (n=6) | 1.498±0.073 (n=6) | 2.02±0.24 (n=6) | 75.90±2.06 (n=6) |
| *Trpm4*<sup>-/-</sup> | ♂ | -643±31 (n=5) | 1.496±0.062 (n=5) | 1.67±0.15 (n=5) | 70.38±1.82 (n=5) |
| control | ♀ | -605±46 (n=4) | 1.327±0.011 (n=4) | 1.48±0.22 (n=4) | 72.38±0.33 (n=4) |
| *Trpm4*<sup>-/-</sup> | ♀ | -584±21 (n=4) | 1.234±0.106 (n=4) | 1.37±0.10 (n=4) | 74.59±2.15 (n=4) |
| **Adult** |  |  |  |  |  |
| control | ♂ | -718±15 (n=10) | 1.437±0.085 (n=10) | 2.07±0.08 (n=10) | 71.13±0.33 (n=10) |
| *Trpm4*<sup>-/-</sup> | ♂ | -605±27 (n=10) | 1.411±0.036 (n=10) | 1.49±0.13 (n=10) | 71.09±0.40 (n=10) |
| control | ♀ | -591±17 (n=8) | 1.320±0.052 (n=8) | 1.41±0.08 (n=8) | 71.22±0.62 (n=8) |
| *Trpm4*<sup>-/-</sup> | ♀ | -562±12 (n=10) | 1.445±0.054 (n=10) | 1.27±0.05 (n=10) | 70.21±0.52 (n=10) |

### Effects of new TRPM4 inhibitors on murine cardiac function

We used the heart explanted-Langendorff system to perform pseudo- and intracardiac ECG recordings in the absence and presence of NBA, a promising, recently described TRPM4 inhibitor (Arullampalam, Preti, Ross-Kaschitza, Lochner, Rougier and Abriel 2021, Ozhathil, Delalande, Bianchi, Nemeth, Kappel, Thomet, Ross-Kaschitza, Simonin, Rubin, Gertsch, Lochner, Peinelt, Reymond and Abriel 2018). Figure 6A shows representative intracardiac ECG traces recorded using control male mice. The A and V waves correspond to electrical activation of the right atria and right ventricle, respectively, and are recorded at two different points of the octopolar probe (Ch5 and Ch1, respectively) (Figure 6A). The A-A and V-V intervals correspond to the time between two electrical activities of the right atrium and the right ventricle, respectively (Figure 6A). The interval between A and V waves (A-V interval) reflects the time required for the electrical impulse to propagate from the right atrium to the right ventricle through the atrioventricular node. Using A-A, V-V, and A-V intervals as readouts for cardiac conduction, we investigated the effect of NBA on male 12-weeks-old *Trpm4* ^-/-^ and control mouse hearts. For this investigation of the potential role of NBA on the heart, young female animals were omitted based on their absence of electrical phenotype. As a control experiment, we investigated the potential effect of the vehicle (DMSO 0.05%) on the A-V interval (Supplementary figure 2). As expected, no effect was observed. Perfusion of 5 µM NBA on young wild-type mouse hearts significantly increased the A-V interval (+44.0% ± 5.1%; n = 6; figure 6B). Unexpectedly, the A-V interval in young male *Trpm4* ^-/-^ mouse hearts was also slightly but significantly increased upon NBA application (+12.4% ± 3.1%; n = 6; figure 6C). Five µM CBA, which is known to inhibit human TRPM4 but not mouse TRPM4 in a heterologous expression system, on the other hand, did not modify the A-V interval (Figure 6B) (Arullampalam, Preti, Ross-Kaschitza, Lochner, Rougier and Abriel 2021). Investigating the A-A and V-V intervals during NBA perfusion on young male wild-type hearts revealed that NBA application led to a third-degree atrioventricular block in 40% of cases. In contrast, none was observed in *Trpm4* ^-/-^ young male hearts. These findings suggest that the NBA-mediated A-V conduction slowing depends mainly, but not completely, on TRPM4 (Figure 6D). Overall, these data demonstrate that the new TRPM4 inhibitor NBA decreases the atrioventricular TRPM4-dependent conduction, which may eventually lead to a third-degree atrioventricular block.

### Effects of new TRPM4 inhibitors on the voltage-gated sodium current

The slight but significant effect of NBA on the A-V interval in *Trpm4* ^-/-^ mouse hearts (figure 6C) suggests that NBA, at 5 µM, may target (an)other element(s) than the TRPM4 channel (Figure 6C). A recent study using canine left ventricle cardiomyocytes suggested that 10 µM CBA can also inhibit the cardiac voltage-gated sodium channel Na_v_1.5 (Dienes et al. 2021). As Na_v_1.5 plays a key role in cardiac electrical impulse propagation, we investigated whether NBA and CBA at the concentration used on isolated murine hearts affected mouse Na_v_1.5, heterologously expressed in HEK-293 cells. No effect was noticed with the vehicle or with 5 µM CBA, but a significant decrease of the sodium current was observed after the application of 5 µM NBA (−33.6 % ± 6.8 %; n = 8) suggesting that NBA, at high concentrations, may inhibit the voltage-gated sodium channel Na_v_1.5 and account for the increase of the A-V interval shown in figure 6C (Figure 7A).

**Figure 7.**
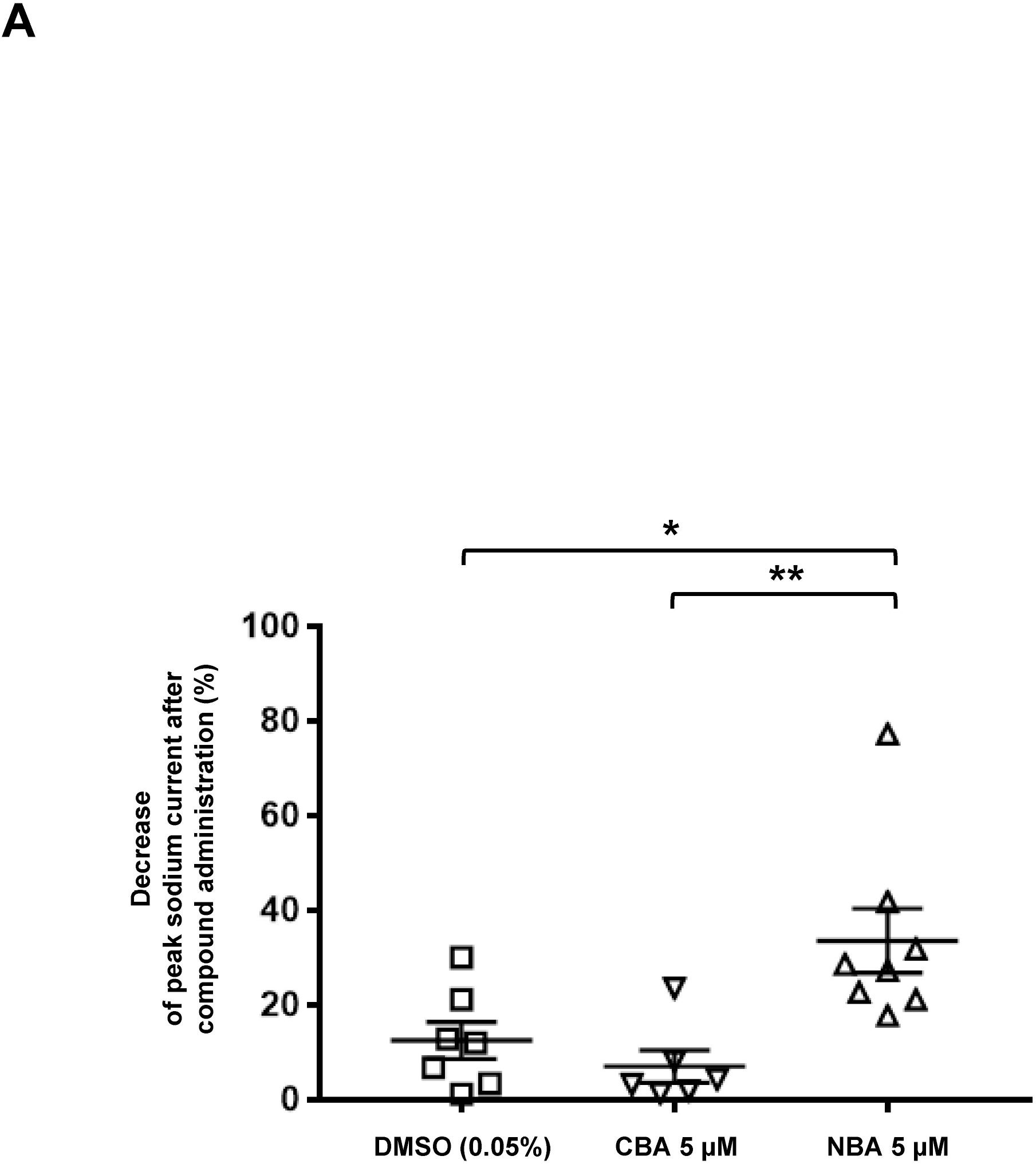
Effects of NBA and CBA on voltage-gated sodium current mediated by mouse Na_v_1.5 in HEK-293 cells. *A*: Dot blots showing the percentage of decrease of sodium current after the application of either the vehicle (DMSO 0.05%), 5 µM CBA, or 5 µM NBA *: p⩽0.05 and **: p⩽0.01 (n⩾6 per group).

## DISCUSSION

Clinical studies have reported a strong link between TRPM4 genetic variants and cardiac conduction disorders (Kecskes et al. 2015, Liu, Chatel, Simard, Syam, Salle, Probst, Morel, Millat, Lopez, Abriel, Schott, Guinamard and Bouvagnet 2013, Palladino et al. 2022, Stallmeyer, Zumhagen, Denjoy, Duthoit, Hebert, Ferrer, Maugenre, Schmitz, Kirchhefer, Schulze-Bahr and Guicheney 2012). Experimental studies have revealed similar findings in mouse models (Demion, Thireau, Gueffier, Finan, Khoueiry, Cassan, Serafini, Aimond, Granier, Pasquie, Launay and Richard 2014, Guinamard et al. 2006, Liu, El Zein, Kruse, Guinamard, Beckmann, Bozio, Kurtbay, Megarbane, Ohmert, Blaysat, Villain, Pongs and Bouvagnet 2010, Mathar et al. 2014, Mathar, Vennekens, Meissner, Kees, Van der Mieren, Camacho Londono, Uhl, Voets, Hummel, van den Bergh, Herijgers, Nilius, Flockerzi, Schweda and Freichel 2010, Pironet et al. 2019). Two recent studies used different *Trpm4* ^-/-^ mouse models to investigate the effect of *Trpm4* gene deletion on the function of the heart (Demion, Thireau, Gueffier, Finan, Khoueiry, Cassan, Serafini, Aimond, Granier, Pasquie, Launay and Richard 2014, Mathar, Kecskes, Van der Mieren, Jacobs, Camacho Londono, Uhl, Flockerzi, Voets, Freichel, Nilius, Herijgers and Vennekens 2014). At basal conditions, Demion *et al*. found that *Trpm4* ^-/-^ mice display multi-level conduction delays and arrhythmogenic activity (Demion, Thireau, Gueffier, Finan, Khoueiry, Cassan, Serafini, Aimond, Granier, Pasquie, Launay and Richard 2014). On the other hand, Mathar and colleagues found that *Trpm4* ^-/-^ hearts exhibited greater contractility in response to β-adrenergic stimulation than controls (Mathar, Kecskes, Van der Mieren, Jacobs, Camacho Londono, Uhl, Flockerzi, Voets, Freichel, Nilius, Herijgers and Vennekens 2014), but they did not observe major cardiac alterations without any stress (Mathar, Kecskes, Van der Mieren, Jacobs, Camacho Londono, Uhl, Flockerzi, Voets, Freichel, Nilius, Herijgers and Vennekens 2014). The discrepancy observed between this two different *Trpm4* ^-/-^ mouse models in basal conditions has been proposed to be due to the different genetic backgrounds (129/SvJ for Mathar *et al*. and C57Bl/6J for Demion *et al*.) (Medert, Pironet, Bacmeister, Segin, Londono, Vennekens and Freichel 2020). Based on these observations, we generated a third *Trpm4* ^-/-^ mouse strain on a C57BL/6JRj background to increase our understanding of the role of TRPM4 under different genetic backgrounds.

Basic phenotyping characterization revealed that the body weight was 9.0% lower in adult male *Trpm4* ^-/-^ than in adult wild-type animals. Surprisingly, Demion and colleagues noticed the opposite effect on body weight in adult male *Trpm4*^*-/-*^ (male wild-type animals had lower body weight than *Trpm4* ^-/-^ mice) (Demion, Thireau, Gueffier, Finan, Khoueiry, Cassan, Serafini, Aimond, Granier, Pasquie, Launay and Richard 2014). However, in addition to the strain of the animal, the substrain of the mice can also lead to drastic differences in terms of metabolic response, which may explain such observations (Ge, Yeung, Mak and Ip 2019, Siersbaek, Ditzel, Hejbol, Praestholm, Markussen, Avolio, Li, Lehtonen, Hansen, Schroder, Krych, Mandrup, Langhorn, Bollen and Grontved 2020). Demion *et al*. did not provide further information concerning the substrain of the C57Bl6/J mice. The observed discrepancies and the potential role of the substrain in the effects observed in the *Trpm4* knockdown can thus not be further explained.

No major surface electrocardiogram alteration has been observed during the 12 first weeks of development for male or female animals comparing wild-type and *Trpm4* ^-/-^ mice. At adult age, only male *Trpm4* ^-/-^ showed a decrease in the heart rate and an increase in the HRV. These findings suggest that the cardiac electrical activity is mainly altered in adult male *Trpm4* ^-/-^ animals. The observation that the knockdown of *Trpm4* did not affect the PR and QRS durations is not in line with what has been reported by Demion and colleagues and may be due to differences in how these parameters were measured (Demion, Thireau, Gueffier, Finan, Khoueiry, Cassan, Serafini, Aimond, Granier, Pasquie, Launay and Richard 2014). In addition, the mice’s genetic background (substrain) might have played a role. The fact that female *Trpm4* ^-/-^ mice did not display any cardiac alterations may indicate that the onset of cardiac electrical dysfunction may occur later in life in females than in male *Trpm4* ^-/-^ mice. In addition, sex hormones might play a role in this discrepancy, as already suggested in previous publications, by protecting the female mice from cardiac disorders (Eckstein et al. 2020). It is worth noting that the prevalence of cardiac disorders linked to TRPM4 dysfunction (e.g., Brugada syndrome and atrioventricular block) has been suggested to be more prevalent in males than in females, as observed in this study (Ehdaie et al. 2018).

Echocardiography scans were performed to investigate the potential cardiac structural and hemodynamic alterations due to the deletion of the *Trpm4* gene. A slight increase in the left ventricle mass of male adult *Trpm4* ^-/-^ mice suggested left ventricle hypertrophy and potential dysfunction of the mitral valves. However, no systolic dysfunctions have been observed in adult male and female animals. Interestingly, the parameters suggesting a mitral valve dysfunction were altered in young male *Trpm4* ^-/-^, but not in young female *Trpm4* ^-/-^. Like the observations from the ECG experiments, these findings also suggest that the onset of the cardiac alterations may be delayed in female animals for a yet-unknown reason. One potential mechanism may be that the diastolic dysfunction related to the left ventricular hypertrophy will cause mitral valve stenosis, which causes a vicious circle and leads to poor diastolic function (Klein and Carroll 2006). We did not investigate if this hypertrophy may be related to hyperplasia, as observed by Demion and colleagues, using another *Trpm4* ^-/-^ mouse model (Demion, Thireau, Gueffier, Finan, Khoueiry, Cassan, Serafini, Aimond, Granier, Pasquie, Launay and Richard 2014). To further analyze left ventricular hypertrophy, histological sections of the hearts and ELISAs of cardiac biomarkers such as ANP and/or BNP may be relevant to characterize better the effect of *Trpm4* deletion on left ventricle structure/function. Regarding the potential mitral valve stenosis, histological section of the mitral valve and the lung should be performed to confirm the proposed hypothesis. This study assessed the “right ventricular” systolic function of *Trpm4*^-/-^ mice for the first time. Only male *Trpm4*^-/-^ mice at adult age showed significant alterations in the right ventricle systolic function, but these changes were not seen in their female counterparts. When considering the etiology of this dysfunction, we consider two possibilities. The systolic dysfunction may be due to a direct effect of the *Trpm4* gene deletion, or the gene deletion may cause an indirect impact on right ventricular function by affecting the mitral valve, leading to stenosis, which increases pulmonary artery pressure. Presently, we do not have evidence to discard one of these two possibilities.

Interestingly, intracardiac recordings using the new potent TRPM4 inhibitor NBA suggest that TRPM4 inhibition leads to atrioventricular conduction slowing and/or atrioventricular block in wild-type hearts. Therefore, we propose that, in addition to the TRPM4-dependent effects observed at the ventricular level, this channel may also play a key role in the atrioventricular node. It is worth noting that the *in-vivo* surface ECGs using male *Trpm4* ^-/-^ mice did not show any alteration of the atrioventricular conduction (time between the end of the P wave and the beginning of the QRS complex). However, this interval reflects the time required for the electrical stimulus to travel through all the atria and ventricles. In the intracardiac ECG, the recording leads have been localized only in the “right heart,” while the electrode recording the electrical activity of the “right atrium” is close to the atrioventricular node. This difference may explain the noted discrepancy. Finally, the intracardiac ECG using the NBA compounds (acute inhibition of TRPM4) does not perfectly resemble the situation with the *Trpm4* ^-/-^ mouse model, in which TRPM4 is not inhibited but is absent from conception. The constitutive TRPM4 knockdown may lead to partial compensatory effects explaining these different observations. Nevertheless, these data strongly support the previous observations done in heterologous expression systems suggesting that NBA may be a new reference compound for investigations of the role of TRPM4 in the heart (Arullampalam, Preti, Ross-Kaschitza, Lochner, Rougier and Abriel 2021). Figures 6 and 7, however, show that 5 µM NBA may also have off-target effects. Given that the NBA concentration to inhibit 50% of mouse TRPM4 in a heterologous expression system is twenty times lower (NBA-IC_50-TRPM4_ =0.215 µM), it will be interesting to perform similar experiments with lower doses of NBA to more specifically target TRPM4 (Arullampalam, Preti, Ross-Kaschitza, Lochner, Rougier and Abriel 2021).

### LIMITATIONS

Among the limitations of this study, it is worth noting that TRPM4 is also widely expressed in different organs and tissues, including the brain and the endocrine system. The cardiac dysfunctions observed after constitutive *Trpm4* gene deletion may be due to indirect mechanisms or related to a compensatory mechanism. Similar experiments using inducible *Trpm4* ^-/-^ and/or cardiac-specific knockdown mouse models may be considered to better characterize the role of TRPM4 in cardiac function. It has proven challenging to reproduce data using similar knockdown models, as the two previous publications using different *Trpm4* ^-/-^ mouse models have already been highlighted. The strain and substrain hypotheses are relevant but may not explain all the observed discrepancies. The different techniques used are complex to handle and may yield experimenter-dependent results, too. Overall, our data and the data from Mathar and Demion suggest that the different research groups should collaborate closely in setting up and standardizing those approaches to achieve reproducible results regarding the role of TRPM4 in cardiac tissue. (Demion, Thireau, Gueffier, Finan, Khoueiry, Cassan, Serafini, Aimond, Granier, Pasquie, Launay and Richard 2014, Mathar, Kecskes, Van der Mieren, Jacobs, Camacho Londono, Uhl, Flockerzi, Voets, Freichel, Nilius, Herijgers and Vennekens 2014).

## CONCLUSIONS

In conclusion, the cardiac investigation of this new *Trpm4* ^-/-^ mouse model confirms the important role of TRPM4 in the proper structure and electrical function of the heart. This study also reveals significant differences between male and female animals that have never been reported before. In addition, the investigation of the effects of NBA on heart function highlights the role of TRPM4 in the atrioventricular node and provides the first evidence showing the efficacy of this compound on native tissues such as the heart. However, the discrepancy between the results presented in this study with those from previous publications suggests that the investigation of the role of TRPM4 in cardiac physiology may be influenced by many unidentified factors, such as mouse substrains and hormones.

## FOOTNOTES

We would like to thank the company PolyGene (PolyGene AG, Rümlang, Switzerland) for the *Trpm4* knockdown targeting strategy and the generation of the *Trpm4* knockdown mouse strain and the PD. Dr. Martin Lochner and his group (Institute of Biochemistry and Molecular Medicine Bern, Switzerland) to produce the compounds NBA and CBA.

## Author contributions

PA, MCE, and JSR conceived and designed the experiments. PA, MCE, and JSR collected, analyzed, and interpreted the data. PA, JSR and HA drafted the manuscript.

