## Supplemental information for "Knockdown of the TRPM4 channel alters cardiac electrophysiology and hemodynamics in a sex- and age-dependent manner in mice"

### SUPPLEMENTARY FIGURE LEGENDS

**Supplementary figure 1. Generation of *Trpm4* $^{-/-}$ mouse strain.** *A*: Schematic drawing of the targeted *Trpm4* locus. Two alleles of the *Trpm4* locus are depicted: the wild-type locus (WT) and the targeted allele (TG). After homologous recombination, three lox P sites (blue arrows) are inserted into the genome flanking exon 10 of *Trpm4* (red arrows). Additionally, an FRT-flanked neomycin(green) / lacZ (blue) cassette is inserted upstream of exon 10. The *Trpm4* $^{-/-}$ strain was backcrossed on a C57BL6/JRj background.

**Supplementary figure 2. Effect of DMSO on cardiac electrical activity from young male hearts using intracardiac recordings.** *A*: Dot blots showing the evolution of the A-V interval after application of vehicle (DMSO 0.05%) on wild-type mouse hearts (*A*) (n = 4 per group).
