## Supplementary figures and images for "Knockdown of the TRPM4 channel alters cardiac electrophysiology and hemodynamics in a sex- and age-dependent manner in mice"

### Supplemental figure 1

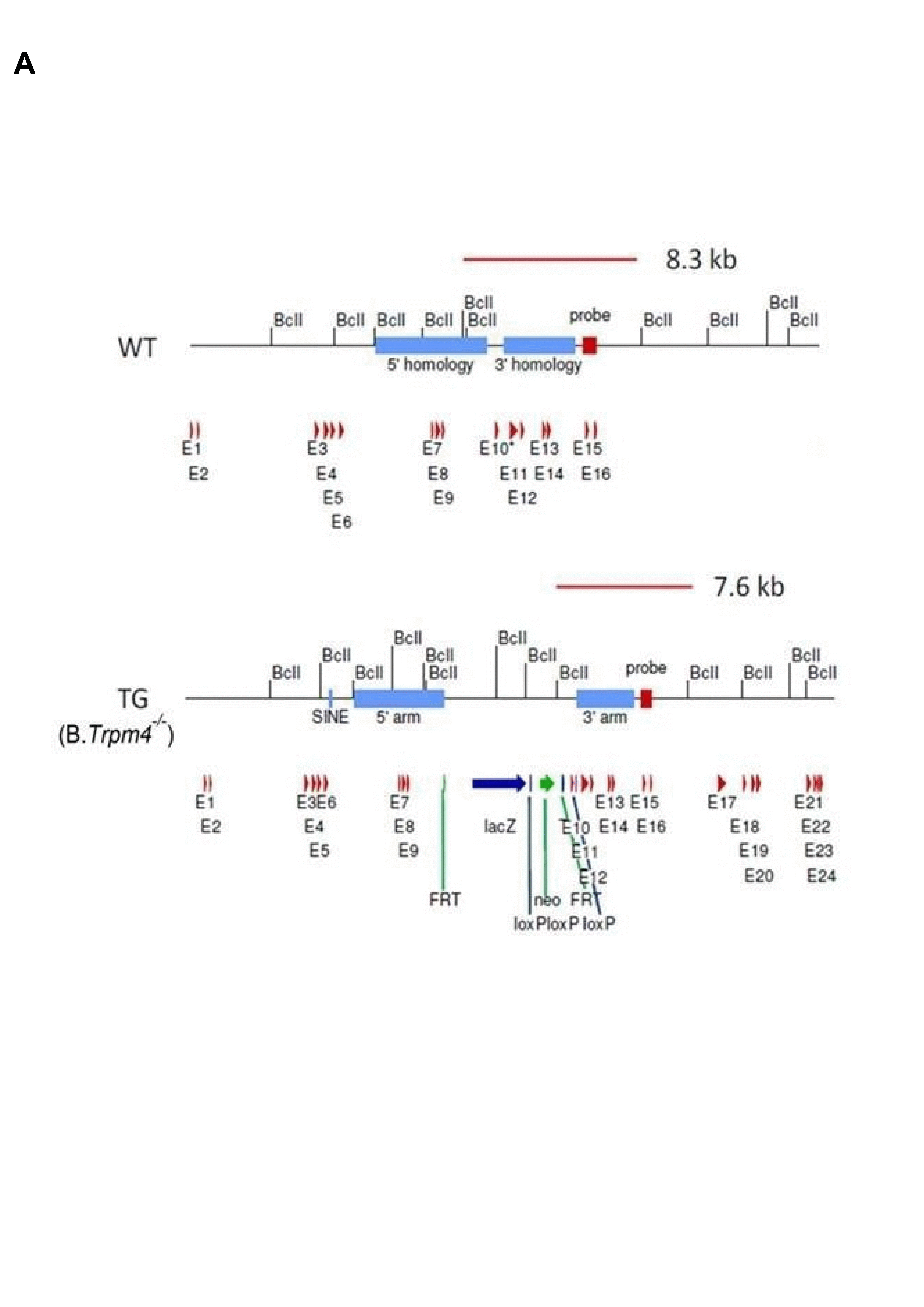

### Supplemental figure 2

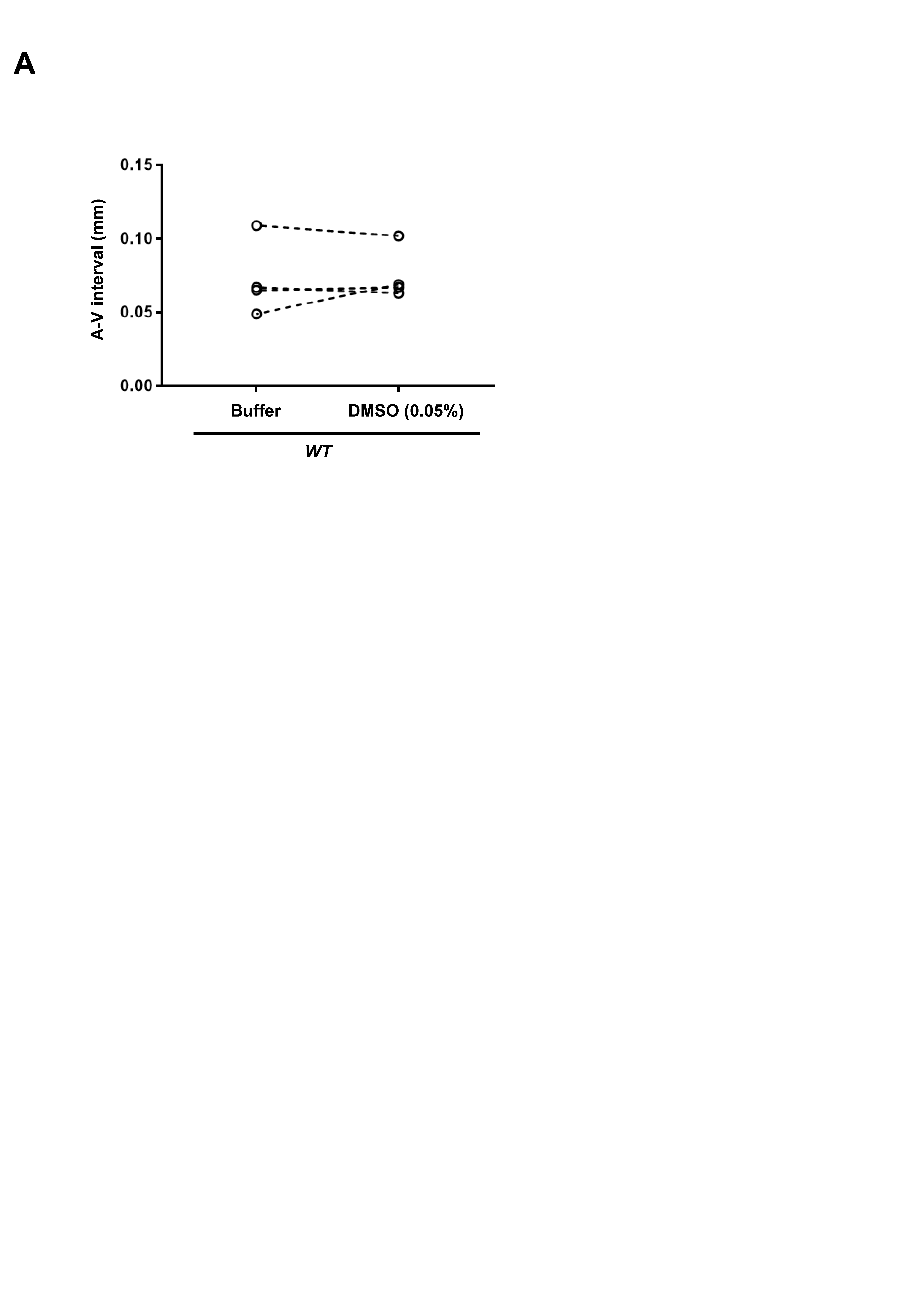
